# MONICA: A Web Application for Automated Whole Optic Nerve Contour Extraction and Morphometric Analysis Validated Across Taxonomic Orders and Image Quality Levels

**DOI:** 10.64898/2026.02.27.707453

**Authors:** Benton Chuter, William White, XiangDi Wang, Linan Guan, Qays Aljabi, Mohamed Moustafa Ibrahim, Lu Lu, Robert W. Williams, TJ Hollingsworth, Monica M. Jablonski

**Affiliations:** Department of Ophthalmology, The University of Tennessee Health Science Center, Memphis, TN, United States; Department of Genetics, Genomics and Informatics, The University of Tennessee Health Science Center, Memphis, TN, United States; Department of Anatomy and Neurobiology, The University of Tennessee Health Science Center, Memphis, TN, United States; The University of South Alabama College of Medicine, Mobile, AL, United States

**Keywords:** optic nerve morphometry, contour extraction, deep learning segmentation, axon density, glial coverage, glaucoma, AxonDeepSeg

## Abstract

Quantitative assessment of optic nerve health requires metrics beyond axon counts alone. Axon density and glial coverage fraction correlate with clinical measures of visual function, yet no existing automated tool extracts optic nerve cross-sectional boundaries to enable their calculation. We developed MONICA (Morphometrics from Optic Nerve Imaging Contour Analysis), a web application that integrates AxonDeepSeg deep learning segmentation with a novel morphology-based contour extraction algorithm to automatically derive whole nerve boundaries alongside axon and myelin masks. The contour extraction algorithm was validated against manual ground truth annotations using 15 optic nerve cross-sections spanning two taxonomic orders (mouse, rabbit), two mouse strains (BXD29, BXD51), and varying preparation quality levels (modern and archival samples). Automated contour extraction demonstrated excellent agreement with manual annotations, achieving an overall Dice similarity coefficient (a measure of segmentation overlap) of 0.987 ± 0.009. Balanced precision (0.985) and recall (0.989) values indicated that the algorithm neither systematically over-segments nor under-segments nerve boundaries. MONICA requires no local software installation and runs entirely in-browser, providing batch processing for high-throughput phenotyping alongside a full suite of per-axon morphometrics. MONICA provides researchers with an accessible tool for complete nerve cross-section morphometry.

## INTRODUCTION

Glaucoma is the leading cause of irreversible blindness worldwide, affecting approximately 64 million people and projected to reach 112 million by 2040 [1]. The disease is characterized by progressive degeneration and loss of retinal ganglion cells (RGCs), with axonal injury in the optic nerve serving as a pathological hallmark [2,3]. Quantitative assessment of optic nerve health through histological analysis of nerve cross-sections enables empirical insight into disease mechanisms and evaluation of neuroprotective interventions. However, total axon counts alone offer an incomplete assessment of nerve health. Axon density, which is influenced by variations in nerve cross-sectional area, provides a more standardized measure for between-subject comparisons [4,5]. Similarly, given the recognized role of astrocyte activation and glial remodeling in glaucomatous neurodegeneration [6,7], characterization of non-axonal glial coverage has emerged as increasingly relevant.

Several automated and semi-automated approaches have been developed for optic nerve morphometry. AxonDeepSeg employs convolutional neural networks to perform three-class semantic segmentation of microscopy images, classifying pixels as axon, myelin, or background with high accuracy [8]. AxoNet 2.0 uses deep learning to count normal-appearing RGC axons and quantify individual axon morphometry from light micrographs [4]. AxonJ offers fully automated, parameter-free axon counting for paraphenylenediamine (PPD)-stained rodent sections [9]. While these tools have substantially advanced the field, they share a common limitation: their primary focus on axon enumeration without concurrent segmentation of optic nerve cross-sectional boundary. This precludes direct calculation of axon density and leaves glial coverage uncharacterized. Further, these tools also have high barriers to entry that preclude their widespread adoption by researchers without computational backgrounds, often requiring local software installation, command-line proficiency or programming experience, and in some cases dedicated GPU hardware. None of them offer an easily accessible browser-based interface.

The absence of automated extraction of the optic nerve cross-sectional boundary is a longstanding limitation in the field of automated quantitative nerve cross-section analysis. Without identification of the pial boundary that defines the nerve perimeter, calculation of true axon density (axons per unit area) requires separate manual measurement of nerve area, introducing additional labor and measurement inconsistency. Likewise, glial coverage, encompassing non-axonal tissue including astrocytes and other glial elements, cannot be quantified without first establishing the outer nerve boundary. Recent evidence suggests that contour-derived metrics such as axon density and glial coverage area ratio may have greater clinical relevance than simple axon counts: in glaucoma animal models, some evidence suggests glial coverage fraction and axon density may correlate more strongly with clinical measures of visual function and intraocular pressure than total axon number [10].

To overcome this limitation, we developed MONICA (Morphometrics from Optic Nerve Imaging Contour Analysis), a web application that integrates the AxonDeepSeg bright-field model with a novel morphology-based contour extraction algorithm. MONICA derives the optic nerve cross-sectional boundary directly from axon and myelin segmentation masks, enabling simultaneous calculation of cross-sectional area, axon density, and glial coverage within a unified analytical workflow. These comprehensive morphometric features enabled by MONICA align with increasing recognition that glaucomatous neurodegeneration extends beyond axon loss alone, as glial activation and tissue remodeling have been identified as key components in optic nerve pathology [6,7].

The primary aim of this study was to validate the contour extraction performance of MONICA against manual ground truth annotations across multiple taxonomic orders, mouse strains, and cross-section image quality levels (encompassing specimen preservation, slide preparation, and imaging conditions). Our secondary aim was to demonstrate that automated extraction of the optic nerve boundary allows for reliable calculation of axon density and glial coverage fraction. We hypothesized that 1) MONICA’s generated contours would demonstrate excellent agreement with manual contour delineation regardless of tissue source or image characteristics, and 2) boundary extraction would allow for accurate and reproducible morphometric measurements, establishing it as a reliable, accessible tool for complete optic nerve morphometry. Our data demonstrate that contour extraction with MONICA allows for accurate and reproducible optic nerve morphometry. This pipeline provides a standardized framework for scalable cross-laboratory implementation.

## RESULTS

### Contour Extraction Performance: Overall

The MONICA contour extraction algorithm demonstrated excellent agreement with manual ground truth annotations across all 15 validation samples (Figure 1A, Table 1). The overall mean Dice coefficient was 0.987 ± 0.009, with individual sample values ranging from 0.967 to 0.996. Intersection over Union (IoU) averaged 0.975 ± 0.017. Precision (0.985 ± 0.017) and recall (0.989 ± 0.011) were well-balanced, indicating that the algorithm neither systematically over-segments nor under-segments the nerve boundary relative to manual annotations.

**Table 1.** Contour extraction validation metrics by tissue group. Values presented as mean ± SD. Dice, Dice similarity coefficient; IoU, Intersection over Union. All metrics demonstrate excellent agreement (>0.96) between automated contour extraction and manual ground truth annotations across species and image quality levels.

| Tissue Group | Dice | IoU | Precision | Recall | F1 Score |
| --- | --- | --- | --- | --- | --- |
| Archival BXD51 (n=5) | 0.987 ± 0.009 | 0.974 ± 0.017 | 0.980 ± 0.021 | 0.993 ± 0.005 | 0.987 ± 0.009 |
| Contemporary BXD29 (n=5) | 0.981 ± 0.009 | 0.962 ± 0.017 | 0.969 ± 0.010 | 0.993 ± 0.007 | 0.981 ± 0.009 |
| Dutch Belted Rabbit (n=5) | 0.994 ± 0.003 | 0.988 ± 0.005 | 0.997 ± 0.003 | 0.984 ± 0.019 | 0.994 ± 0.003 |
| **Overall (n=15)** | **0.987 ± 0.009** | **0.975 ± 0.017** | **0.985 ± 0.017** | **0.989 ± 0.011** | **0.987 ± 0.009** |

**Figure 1.**
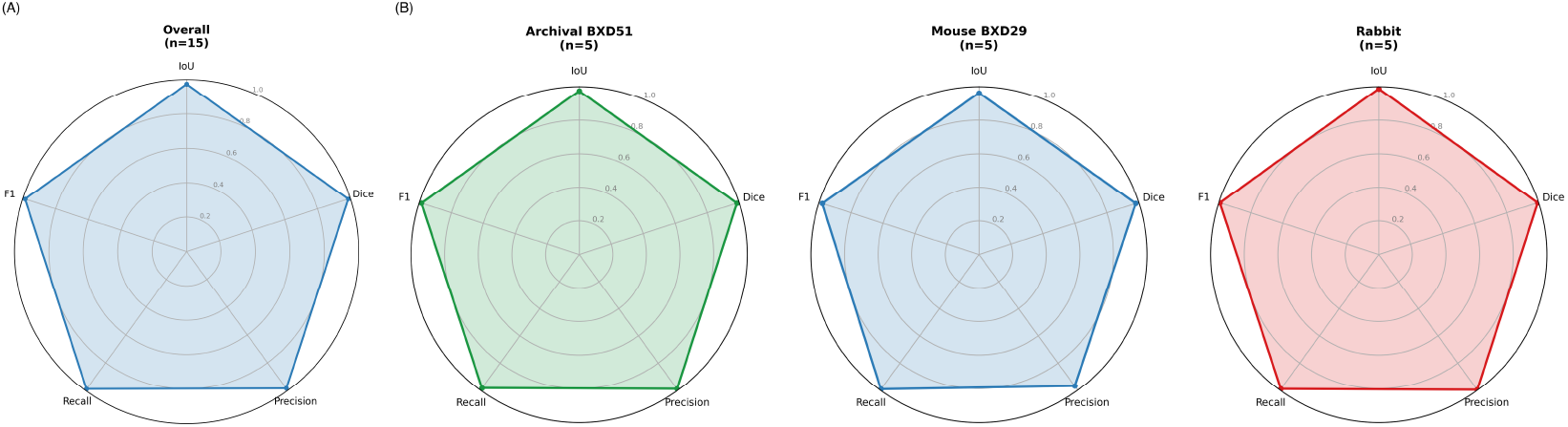
Contour extraction validation performance. Radar plots displaying Dice coefficient, Intersection over Union (IoU), Precision, Recall, and F1 Score for automated contour extraction compared to manual ground truth annotations. (A) Overall performance across all 15 validation samples (mean ± SD): Dice = 0.987 ± 0.009, IoU = 0.975 ± 0.017, Precision = 0.985 ± 0.017, Recall = 0.989 ± 0.011. (B) Performance stratified by tissue group: Archival BXD51 mouse (green, n=5, Dice = 0.987), Contemporary BXD29 mouse (blue, n=5, Dice = 0.981), and Dutch belted rabbit (red, n=5, Dice = 0.994). **Abbreviations:** Dice, Dice similarity coefficient; F1, F1 score; IoU, intersection over union.

### Contour Extraction Performance: Cross-Order and Cross-Quality Validation

Performance was consistent across tissue groups representing different taxonomic orders and preparation quality levels (Figure 1B, Table 1). Rabbit optic nerve samples (n = 5) achieved the highest agreement, with mean Dice = 0.994 ± 0.003 and IoU = 0.988 ± 0.005. Mouse samples from the BXD29 strain (n = 5) achieved Dice = 0.981 ± 0.009 and IoU = 0.962 ± 0.017. Archival mouse samples from the BXD51 strain (n = 5), representing images with variable staining quality and tissue preservation, achieved Dice = 0.987 ± 0.009 and IoU = 0.974 ± 0.017.

The algorithm demonstrated consistent precision across groups (range: 0.969 to 0.997) and recall (range: 0.984 to 0.993), confirming robustness to both taxonomic order differences and preparation quality variation.

### Per-Sample Performance

Individual sample performance metrics are presented in Table 2. All 15 samples achieved Dice coefficients exceeding 0.96, with 12 of 15 samples (80%) exceeding 0.98. The lowest-performing sample (BXD29 M639, Dice = 0.967) still demonstrated excellent agreement with manual annotation. The highest-performing samples were rabbit specimens F243_OD1 (Dice = 0.996) and 021909_78L from the BXD51 archived samples (Dice = 0.995).

**Table 2.** Per-sample contour extraction validation metrics. All 15 samples achieved Dice coefficients exceeding 0.96, with 12 of 15 samples (80%) exceeding 0.98. Dice, Dice similarity coefficient; IoU, Intersection over Union; F1, F1 Score (equivalent to Dice).

| Group | Sample ID | Dice | IoU | Precision | Recall | F1 |
| --- | --- | --- | --- | --- | --- | --- |
| BXD51 | 021909_77L | 0.992 | 0.985 | 0.988 | 0.997 | 0.992 |
| BXD51 | 021909_78L | 0.995 | 0.991 | 1.000 | 0.991 | 0.995 |
| BXD51 | 021909_81L | 0.987 | 0.975 | 0.981 | 0.994 | 0.987 |
| BXD51 | 090809_84L | 0.974 | 0.949 | 0.950 | 0.999 | 0.974 |
| BXD51 | 090809_85L | 0.985 | 0.970 | 0.982 | 0.987 | 0.985 |
| BXD29 | F641 | 0.988 | 0.977 | 0.979 | 0.998 | 0.988 |
| BXD29 | F642 | 0.985 | 0.970 | 0.971 | 0.999 | 0.985 |
| BXD29 | F643 | 0.983 | 0.966 | 0.967 | 0.999 | 0.983 |
| BXD29 | M639 | 0.967 | 0.936 | 0.955 | 0.979 | 0.967 |
| BXD29 | M640 | 0.981 | 0.963 | 0.973 | 0.989 | 0.981 |
| Rabbit | F241_OS2 | 0.996 | 0.992 | 0.999 | 0.993 | 0.996 |
| Rabbit | F243_OD1 | 0.996 | 0.992 | 0.999 | 0.993 | 0.996 |
| Rabbit | F243_OS1 | 0.992 | 0.984 | 0.994 | 0.990 | 0.992 |
| Rabbit | M232_OD1 | 0.995 | 0.990 | 0.999 | 0.991 | 0.995 |
| Rabbit | M232_OS1 | 0.990 | 0.981 | 0.994 | 0.952 | 0.990 |

Visual inspection of segmentation overlays (Figure 2) confirmed that the algorithm successfully identified whole-nerve boundaries across diverse tissue types, including nerves with elliptical cross-sections, variable staining intensity, and edge artifacts.

**Figure 2.**
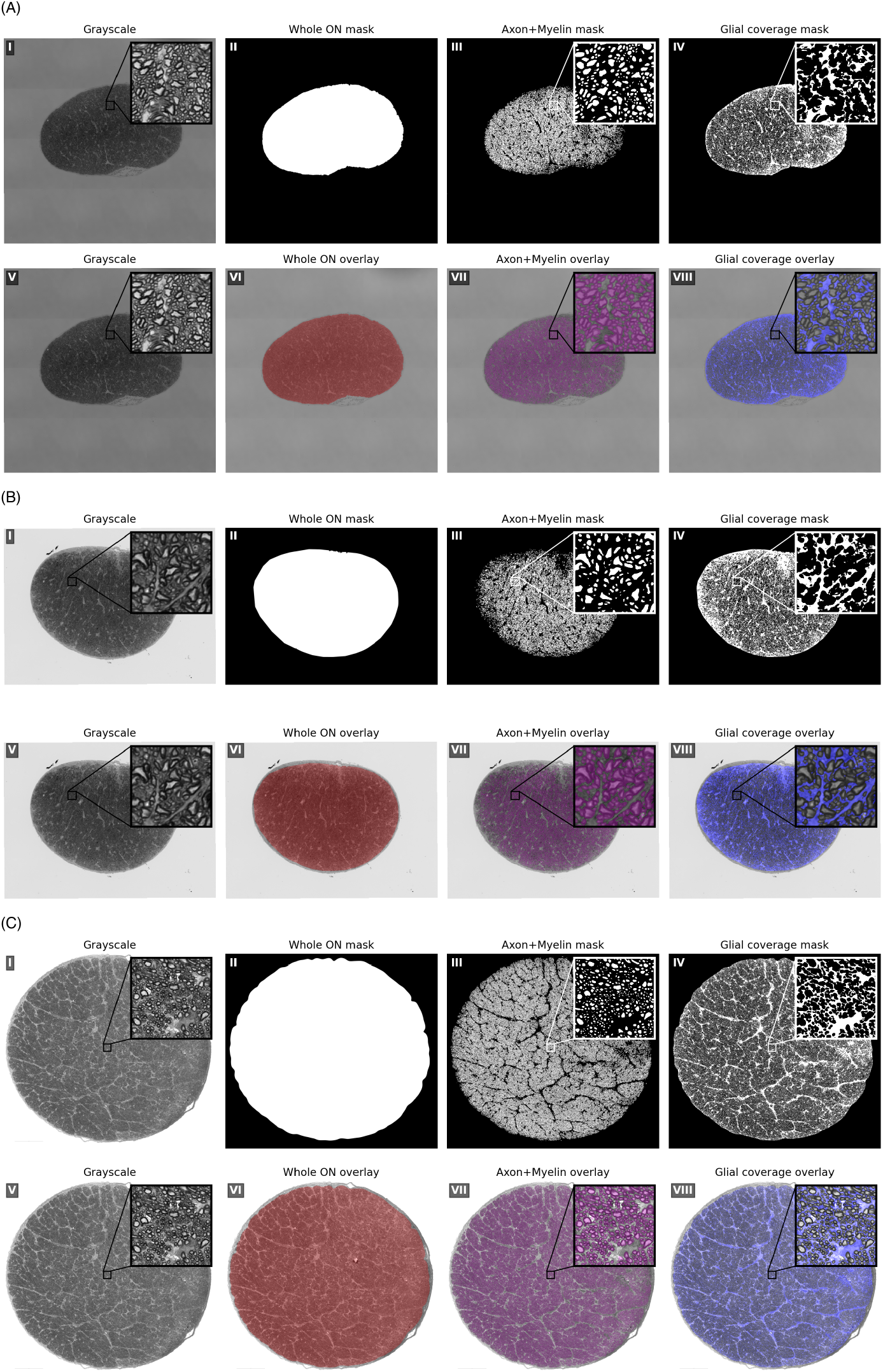
Representative MONICA pipeline outputs across validation groups. Each sample is shown as a 2×4 panel: grayscale input (I, V), whole optic nerve (ON) contour mask and overlay (II, VI), axon+myelin segmentation mask and overlay (III, VII), and glial coverage mask and overlay (IV, VIII). High-magnification insets (10× panel view) highlight axon- and glial-level detail with boxes marking the sampled region. (A) Archival BXD51 mouse (021909_78L). (B) Contemporary BXD29 mouse (F641). (C) Dutch belted rabbit (GP1 M232 OD1).

### Representative Processing Outputs

Figure 2 displays representative MONICA outputs for each validation group. For each sample, the algorithm generated: (1) whole optic nerve contour mask, (2) combined axon and myelin segmentation mask from AxonDeepSeg, and (3) derived glial coverage mask representing non-axonal tissue within the nerve boundary. The contour extraction algorithm successfully captured the pial boundary regardless of nerve size, with rabbit nerves being substantially larger than mouse samples.

## DISCUSSION

This study validated MONICA, a web application for automated whole optic nerve contour extraction and morphometric analysis. The contour extraction algorithm demonstrated excellent agreement with manual ground truth annotations, achieving an overall Dice coefficient of 0.987 across 15 samples representing two taxonomic orders (Rodentia and Lagomorpha), two mouse strains, and varying preparation quality levels. Performance was consistent regardless of tissue source, with all validation groups achieving Dice coefficients exceeding 0.98 on average. These results establish MONICA as a reliable tool for automated whole-nerve boundary detection, answering a longstanding unmet need in existing optic nerve morphometry pipelines.

The primary contribution of this work lies in providing an easily accessible pipeline requiring minimal technical or computational resources that can calculate highly relevant morphometric parameters that existing tools do not provide. Current automated pipelines for nerve cross-section analysis focus on axon enumeration without concurrent segmentation of whole-nerve boundaries [4,8,9]. This limitation necessitates separate manual measurement of nerve cross-sectional area to calculate axon density, introducing additional labor and potential for measurement inconsistency. By deriving the nerve boundary directly from axon and myelin segmentation masks, MONICA provides coincident area measurement that enables direct calculation of axon density within a single automated workflow. The algorithm’s success across all 15 validation samples, including nerves with substantial tissue heterogeneity, confirms its reliability for routine research applications.

The cross-order, cross-strain, and cross-quality validation strategy employed here provides strong evidence for algorithm generalizability. The inclusion of rabbit samples alongside mouse samples demonstrates that the morphology-based approach generalizes across orders (Rodentia and Lagomorpha) with substantially different nerve dimensions, and the algorithm’s success with the larger rabbit optic nerves suggests potential compatibility with human samples. The inclusion of archival samples (BXD51) alongside more recent preparations (BXD29, rabbit) confirms robustness of MONICA to preparation quality variation, an important consideration for laboratories with existing image archives. The consistent performance across these diverse conditions (Dice range: 0.981 to 0.994 for group means) suggests that MONICA can be deployed across varied research contexts without requiring parameter adjustment or retraining. Although validated here on optic nerve, the morphology-based contour extraction approach is not anatomically specific and may generalize to other myelinated nerve cross-sections, including spinal cord white matter tracts and peripheral nerves prepared with similar staining protocols.

The glial coverage area ratio, which cannot be calculated without first establishing the optic nerve cross-sectional boundary, may capture pathological changes in the non-neuronal compartment that simple axon counts miss. Future studies with larger cohorts should investigate whether these contour-derived metrics provide independent prognostic information beyond axon enumeration.

Several limitations warrant consideration. The validation dataset, while diverse in taxonomic order, strain, and preparation quality, comprised samples from a single laboratory. Performance on images from other non-optic nerve sources remains to be established. The glial coverage area calculations assume that all tissue within the nerve boundary not classified as axon or myelin represents glial coverage; more refined classification of subtypes would require additional segmentation capabilities not currently implemented.

MONICA is freely accessible as a web application at monica.jablonskilab.org. The tool accepts standard bright-field microscopy image formats and provides downloadable outputs including segmentation masks, contour boundaries, visualization overlays, and morphometric summary statistics. No local software installation or computational expertise is required, making complete nerve morphometry accessible to researchers regardless of programming background or computational resource access. The batch processing mode enables analysis of entire experimental cohorts in a single session, supporting high-throughput phenotyping workflows common in glaucoma genetics research. For computational researchers, MONICA also provides a programmatic REST API with an accompanying Python client library, enabling scripted analysis pipelines and integration with existing laboratory information management systems.

In conclusion, MONICA provides validated, automated extraction of the optic nerve cross-sectional boundary with excellent agreement to manual annotation across taxonomic orders and preparation quality levels. By enabling calculation of axon density and glial coverage fraction alongside traditional morphometrics, MONICA fills an unmet need in nerve cross-section morphometry tools. MONICA will benefit high-throughput phenotyping in glaucoma research and other contexts requiring standardized nerve characterization.

## METHODS

### Study Design and Datasets

This validation study compared automated contour extraction from the MONICA pipeline against manual ground truth annotations using optic nerve cross-sections from multiple taxonomic orders and preparation quality levels.

Contour Validation Dataset comprised 15 bright-field microscopy images of paraphenylenediamine (PPD)-stained, plastic-embedded optic nerve cross-sections[11] organized into three groups: (1) Archival BXD51 Mouse (n = 5): Archival images prepared in 2009 from the Jablonski Laboratory collection, representing samples with inconsistent preparation, staining, and imaging quality as well as variable tissue preservation characteristic of archival datasets. (2) Contemporary BXD29 Mouse (n = 5): High-quality images processed in 2024, representing optimal contemporary preparations. (3) Dutch belted Rabbit (n = 5): Cross-sectional images from a rabbit glaucoma model prepared in 2023, representing cross-order generalization to larger nerve dimensions.

Images were acquired using a Zeiss Axio Observe equipped with a 63x objective, the image tiling feature was used to stitch multiple individual images together to form a composite image of the entire nerve. Ground truth contour annotations were created by an expert observer (B.C.) who manually delineated the pial boundary of each nerve using polygon annotation tools. Binary masks were generated from annotations for validation analyses.

All animal procedures were conducted in accordance with protocols approved by the Institutional Animal Care and Use Committee (IACUC) of the University of Tennessee Health Science Center and adhered to the ARVO Statement for the Use of Animals in Ophthalmic and Vision Research.

### MONICA Web Application

MONICA (Morphometrics from Optic Nerve Imaging Contour Analysis) is a GPU-accelerated web application that integrates deep learning axon segmentation with automated contour extraction. The application is freely accessible at https://monica.jablonskilab.org and supports both single-image analysis and batch processing of multiple images (Figure 3). The interface accepts standard bright-field microscopy formats (PNG, TIFF) and provides downloadable outputs including segmentation masks, contour boundaries, visualization overlays, and morphometric measurements in CSV format. Processing is performed on server-side GPU hardware, requiring no local software installation or computational resources from the user.

**Figure 3.**
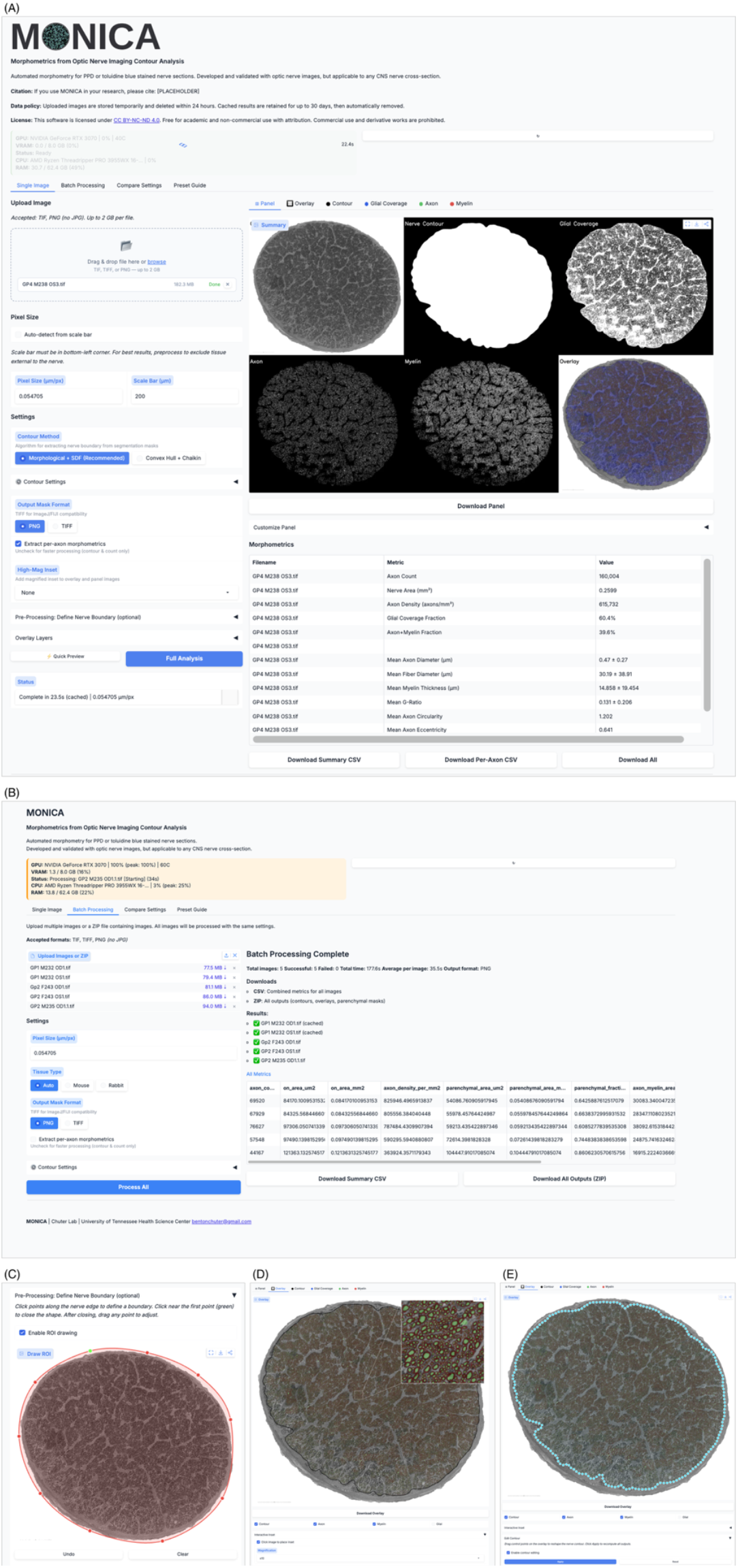
MONICA web application interface. (A) Single image analysis mode allowing users to upload individual nerve cross-section images, adjust processing parameters and region of interest, and view results interactively. (B) Batch processing mode enabling automated analysis of multiple images with progress tracking and consolidated output generation. (C) Region of Interest Selection. (D) Interactive Inset. (E) Contour editing.

The application includes a compare settings mode for side-by-side evaluation of different contour extraction parameters, a quick preview mode that operates on downsampled images for rapid parameter tuning, and five built-in parameter presets (Default, Aggressive, Conservative, Scattered, Dense) calibrated for different tissue preparation qualities. Post-processing visualization features include interactive overlay customization with per-layer toggles for contour, axon, myelin, and glial coverage masks; clickable high-magnification inset placement at user-selected locations (x3, x5, x10, x20); an interactive spline-based contour editor that allows manual refinement of the extracted nerve boundary via 200 draggable control points, with all downstream metrics and masks automatically recomputed upon adjustment; and configurable multi-panel figure generation.

#### Scale Bar Calibration

Pixel-to-micrometer calibration is performed using automated scale bar detection. The algorithm locates the scale bar in standard lower-left position, measures its length in pixels, and computes pixel size based on known physical length. Manual override is available.

#### AxonDeepSeg Segmentation

Axon and myelin segmentation is performed using AxonDeepSeg (version 5.3.0), an open-source deep learning framework for automated segmentation of myelinated axons [8]. The pipeline employs the generalist bright-field model, which generates three-class semantic segmentation masks distinguishing axon (intra-axonal space), myelin sheath, and background regions. Inference is performed using nnU-Net with a cached predictor architecture: the model is loaded once at application startup and reused across all subsequent requests, with test-time augmentation (mirroring) disabled and prediction performed entirely in memory without intermediate file I/O. These optimizations reduce per-image inference latency while maintaining numerically identical segmentation output.

#### Morphology-Based Contour Extraction

Whole-nerve boundaries are derived from combined axon and myelin segmentation masks through the following steps: (1) Mask Consolidation: Binary axon and myelin masks are merged into a unified nerve tissue mask; (2) Morphological Operations: The consolidated mask undergoes morphological closing (rectangular structuring element) to fill gaps between adjacent axon-myelin units, followed by morphological opening to remove isolated pixels; (3) Boundary Detection: The outer boundary of the processed mask is extracted using contour detection algorithms (OpenCV findContours). For images with multiple detected contours, the contour enclosing the largest area is selected; and (4) Contour Smoothing: The detected boundary undergoes spline interpolation to produce anatomically plausible nerve outlines. Following contour extraction, axon and myelin segmentation masks are filtered to exclude components whose centroids fall outside the derived nerve boundary. This step removes spurious segmentation detections in background tissue or embedding artifacts that would otherwise inflate axon counts and bias morphometric calculations.

#### Derived Metrics

MONICA generates two categories of morphometric output. Summary metrics calculated from the extracted contour include: optic nerve cross-sectional area, axon count, axon density (axons per mm^2^), glial coverage, and glial coverage fraction. Per-axon morphometrics include: axon area, equivalent circular diameter, perimeter, circularity, eccentricity, solidity, orientation, minimum and maximum Feret diameters, fiber (axon plus myelin) area and diameter, myelin area, myelin thickness, and g-ratio (ratio of axon diameter to fiber diameter). Aggregate statistics (mean, standard deviation, median, minimum, maximum) are computed for all per-axon measurements. Users may optionally skip detailed per-axon analysis for faster processing when only summary metrics are required.

### Validation and Statistical Analysis

Agreement between automated and manual contour masks was assessed using: Dice Similarity Coefficient (DSC), calculated as 2 × |A ∩ B| / (|A| + |B|), where A and B represent automated and manual binary masks (values range from 0 to 1); Intersection over Union (IoU), calculated as |A ∩ B| / |A ∪B|; Precision, |A ∩ B| / |A|, representing the proportion of automated segmentation that overlaps with ground truth; Recall, |A ∩ B| / |B|, representing the proportion of ground truth captured by automated segmentation; and F1 Score, the harmonic mean of precision and recall.

### Software and Implementation

The web application is deployed on institutional computing infrastructure equipped with an NVIDIA GeForce RTX 3070 (8 GB VRAM), AMD Ryzen Threadripper PRO 3955WX 16-core processor, and 64 GB system memory. The processing pipeline was implemented in Python 3.11 with key dependencies including AxonDeepSeg (v5.3.0), nnU-Net (v2.2.1), PyTorch (v2.3.1), NumPy, SciPy, OpenCV, Pandas, and scikit-image. Public access is provided through a Cloudflare tunnel with chunked file upload support for images up to 2 GB, bypassing standard request body size limits. GPU convolution kernels are auto-tuned via cuDNN benchmarking, and morphometric calculations use vectorized NumPy operations for batch computation of per-axon measurements.

## Code and Data Availability

MONICA is freely accessible at https://monica.jablonskilab.org. The AxonDeepSeg framework is available at https://github.com/neuropoly/axondeepseg. Ground truth datasets are available from the corresponding author upon reasonable request.

## AUTHOR CONTRIBUTIONS

**B.C**.: Conceptualization, Methodology, Validation, Software, Formal analysis, Visualization, Writing – original draft, Writing – review & editing. **W.W**.: Methodology, Resources, Investigation. Writing, review & editing. **XW**: Investigation. **LG**: Investigation. **Q.A**.: Writing – review & editing. **M.M.I**.: Investigation. **L.L**.: Resources, Writing - review & editing. **R.W.W**.: Resources, Writing - review & editing. **T.J.H:** Writing – review & editing. **M.M.J**.: Conceptualization, Supervision, Project administration, Funding acquisition, Writing - review & editing

